# MILD TRAUMATIC BRAIN INJURY IMPAIRS SPATIAL WORKING MEMORY IN RATS

**DOI:** 10.1101/2025.04.28.650881

**Authors:** Gabriel D. Nah, Andrea G. Hohmann, Nicholas Port, Jonathon D. Crystal

## Abstract

Mild Traumatic Brain Injury (mTBI), or concussion, is the most common form of traumatic brain injury, which accounts for about 80% of cases. It is a common problem in contact sports and may lead to cognitive impairment. This study used the Wayne State University closed-head weight-drop model in lightly anesthetized and unrestrained Long Evans rats. This model allows for the rapid acceleration and deceleration of the head and torso, similar to the biomechanics in human mTBI. Rats were administered a single weight drop. Sham animals were treated the same as the mTBI group but were not subjected to weight drop. Rats were trained in an 8-arm radial maze to assess spatial working memory before and after weight drop manipulation. We observed that the injured rats’ spatial working memory performance significantly declined compared to the sham rats (cohen’s d = 1.88). Specifically, the performance of the sham group continued to improve after the sham procedure, whereas the performance of the injury group decreased. This study suggests the WDM model produces a deficit in spatial working memory in rats.

## INTRODUCTION

Over 3 million people are affected by mild traumatic brain injury (mTBI), commonly referred to as concussion, every year in the United States (Cassidy et al., 2004; Maas, Stocchetti, & Bullock, 2008). TBI is a neurological episode whereby head injury actuates molecular, cellular, and structural pathology, which may bring about functional disturbance to the central nervous system (Rehman, Ali, Tawil, & Yonas, 2008). It can also be classified as an impact to the head, resulting in rapid acceleration and deceleration of the head, neck, and torso. TBI exists on a spectrum, from asymptomatic mild injuries to severe TBI, involving high focal contact forces that may produce skull fracture, subdural hematoma, cerebral damage, or hemorrhage within the hemispheres (Bogoslovsky, Gill, Jeromin, Davis, & Diaz-Arrastia, 2016; Hiskens, Angoa-Perez, Schneiders, Vella, & Fenning, 2019). Common causes of TBI include car accidents, sports, and falls, which can lead to temporary or permanent impairments of cognitive and physical functions (Xiong, Mahmood, & Chopp, 2013). Frequent signs and symptoms of mTBI are headaches, light and sound sensitivity, balance issues, disrupted sleep, and memory problems (Brooks et al., 2017).

In previous studies, patients were most concerned with memory impairment after an mTBI (Flynn, 2010). Some experienced loss of consciousness during the impact, post-traumatic amnesia, short-term and long-term memory impairments, and working memory problems (Vakil, 2005). Working memory impairment in humans has been studied in various experiments (Kumar, Rao, Chandramouli, & Pillai, 2009), but the impact of mTBI on spatial working memory still needs to be further studied (Useros Olmo, Perianez, Martinez-Pernia, & Miangolarra Page, 2020). Prior human studies have required participants to perform in a visual-spatial working memory task (Gorman, Barnes, Swank, Prasad, & Ewing-Cobbs, 2012; Kumar et al., 2009). Due to rats’ visual systems not being as robust as humans, researchers rely on other models to assess spatial memory. The Morris Water Maze is a commonly used behavioral method for evaluating spatial working memory. However, studies that have used this method may have used a more moderate injury apparatus, which may have been associated with motor impairment, which may have interfered with the rat’s ability to accomplish the task (Hamm et al., 1996; Smith et al., 2014). To account for the valuable but potentially confounding finding, researchers may use a less severe injury model and a sensitive behavioral model. This study’s objective was to fill this gap in the literature of mild traumatic brain injury.

Several animal injury models of mTBI include controlled cortical impact, fluid percussion injury, and weight drop models (WDM) (Romeu-Mejia, Giza, & Goldman, 2019). These injury models have been shown to produce impairments on a behavioral and neurological level, similar to human data (Qin et al., 2018). However, these models also have limitations in their design that reduce translation from animal models to humans with mTBI. For example, in some studies, the animal’s head is restrained or is in a fixed position, craniotomies are performed to impact the brain more directly, and sometimes, the model has a high mortality rate. Those features could classify those models as moderate or severe injuries rather than mild (Hiskens et al., 2019). Therefore, the Wayne State University Closed-Skull Weight Drop model was created to reduce these limitations, thus making the injury model more human-like (Masse et al., 2019).

Previous studies have used methods like the Morris Water Maze, Y-maze, and the 8-arm radial maze to assess spatial working memory (Chen, Hong, Wang, & Chang, 2021; Darwish, Mahmood, Schallert, Chopp, & Therrien, 2012; Pham et al., 2021). Typically, the 8-arm radial maze assesses spatial working memory by dividing sessions into two phases, referred to as the study-test procedure (Crystal & Babb, 2008). Food is available once at each of the eight locations. The study phase allows rats to obtain food at four randomly selected arms. In the test phase, all arms are accessible, previously inaccessible arms are baited, and previously visited arms are unbaited. Spatial working memory is measured by the degree to which the rat restricts its visits to baited arms in the test phase while avoiding revisits to unbaited arms (Roberts 1998; Olton & Samuelson 1976).

We trained rats to perform a spatial working memory task using the 8-arm radial maze. Then, we used the Wayne State University WDM to determine if the weight drop produces an impairment in spatial working memory relative to sham control. This weight-drop model allows for the unrestrained rat to undergo the biomechanical motion commonly seen in humans. Specifically, the rapid acceleration and deceleration of the head, neck, and torso.

## METHODS

### Animals

Twenty male and female Long Evans rats (Rattus norvengicus; Envigo, Indianapolis, IN) were housed individually and maintained on a 12-hour light/dark cycle with light onset at 7:50 and offset at 19:50 EST. Upon arrival in the lab at the age of 7 weeks, the rats were given unlimited access to 5012-Rat-Diet (PMI Nutrition International, St. Louis, MO) for one week. Next, male rats received 15 g/day, and female rats received 11g/day of food. Water was available ad libitum, except during brief testing sessions. All procedures followed national guidelines and were approved by the Bloomington institutional animal care and use committee at Indiana University. Four rats (2 male, 2 female) did not meet our learning criterion and were excluded from the study.

### Apparatus

An 8-arm radial maze (positioned 81 cm above the floor) consisted of a central hub (29 cm diameter, 11 cm high), guillotine doors, and a food trough and pellet dispenser at the distal end of each arm. Experimental events (movement of guillotine doors, activation of food dispensers, and interruption of photo beams) were controlled by a computer running Windows XP. Data were recorded (10-ms resolution) with MED-PC software (version 4.1). Pellets were placed outside each runway in perforated bags in order to keep food odors constant throughout all parts of the experiment. The maze was cleaned with 2% chlorohexide before placing each rat in the maze.

### Maze Training

Pretraining consisted of maze acclimation in ten sessions (one session per day for each rat). A session started 30 seconds after a rat was placed in the maze, at which point all the doors opened. Each rat had 10 minutes to explore the maze. All arms were baited with five food pellets (one at the door, three in the arm, and one in the food trough). All guillotine doors closed when a session ended, and the rat was removed from the maze. Uneaten pellets were discarded, and the maze was cleaned. Maze acclimation was assessed by how many food pellets were left in the maze after each session.

#### 8-Arm Training

8-arm training consisted of 15 daily sessions (one session per day for each rat). All eight doors opened simultaneously, and each location provided food (Figure 1). A food pellet was dispensed when the photobeam in the food trough was first interrupted, which was considered a correct choice. A revisit to an arm did not produce additional food and was considered an error. The session ended after the rat visited all arms or after 10 minutes had elapsed. 8-arm training performance was assessed by how many correct arm visits were made in a rat’s first eight choices.

**Figure 1.**
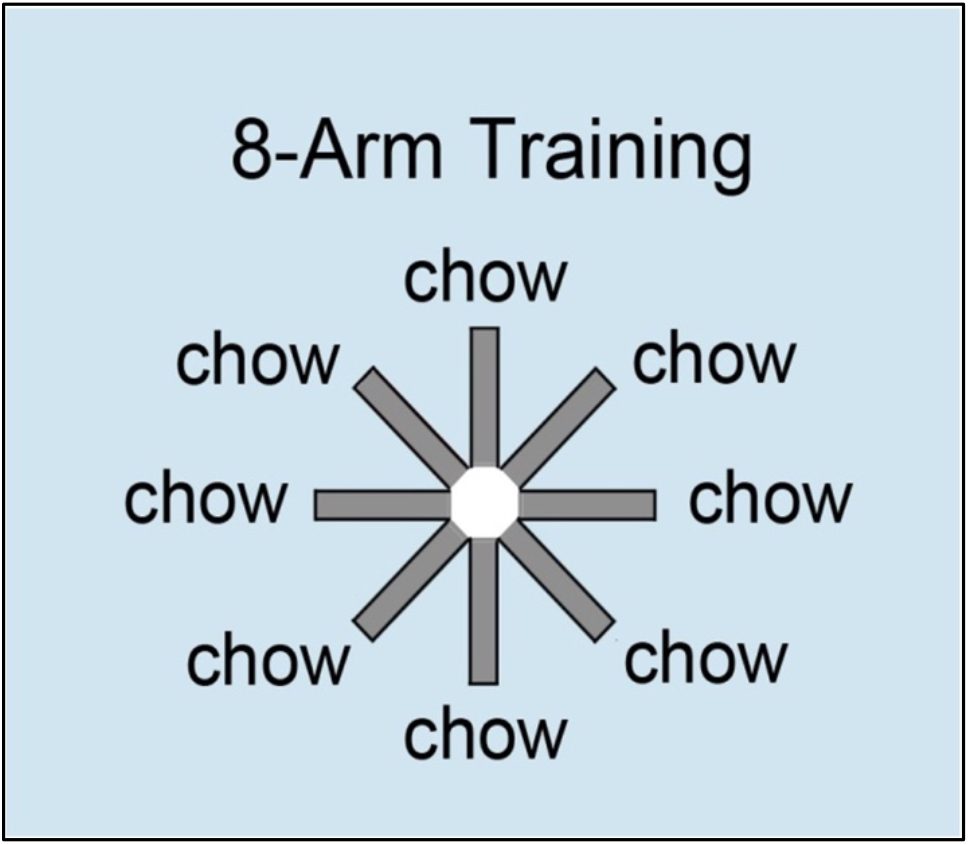
Schematic of the 8-arm radial maze during 8-arm training. Food rewards are located at the end of each runway only once during each daily session. All eight doors are open and accessible to the animal.

### Study-Test Procedure

The Study-Test procedure consisted of two phases: a study phase and a test phase (Figure 2). The study phase began with a rat placed in the central hub. Four randomly selected doors opened 30 seconds after the rat was placed in the hub. Each arm provided a food pellet contingent on the first visit. Revisited arms did not offer a food pellet. The study phase ended when all four accessible food troughs had been visited or after 10 minutes. Next, the rat was removed, the maze was cleaned, and the rat was placed back into the central hub to start the test phase. The test phase began 30 seconds after the program was initiated, at which point all eight maze doors were opened. Food was available in arms that were inaccessible in the test phase. Visits to arms that were inaccessible in the study phase provided a food pellet in the test phase and were considered the correct choices. A revisit did not produce additional food and was considered an error. The test phase ended when the four baited arms were visited or after 10 minutes. Accuracy was defined as the proportion of correct visits in a rat’s first four choices in the test phase. Study-Test sessions were conducted for 20 daily sessions before sham/weight drop (WtDp) induction. Rats were required to meet a criterion of 75% correct in the last five training sessions to advance to weight drop or 8-arm induction. Four rats were excluded from the experiment due to not meeting the criterion. The remaining rats completed 14 sessions after sham/WtDp induction.

**Figure 2.**
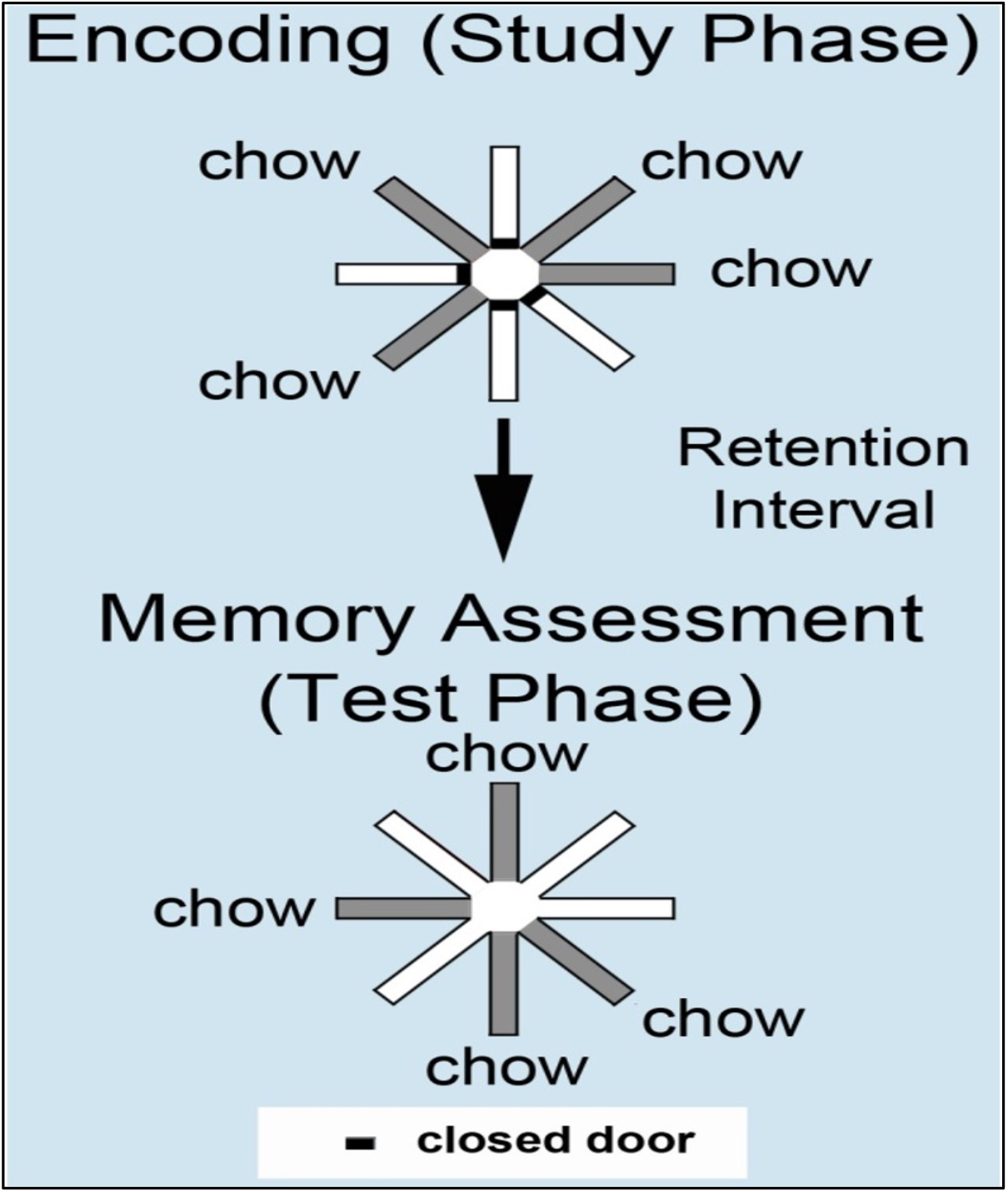
Schematic of the Study-Test procedure. In the study phase, four randomly selected arms are opened to the rat. Rats receive food rewards once from each of the accessible arms. In the test phase, all eight doors are open. Rats receive food rewards once from the previously inaccessible arms from the study phase.

### Mild Traumatic Brain Injury Animal Model

Once baseline data in the study-test phase was obtained, a modified version of the Wayne State University WDM (Figure 3) was used to induce mTBI. Rats were anesthetized using sodium isoflurane. The rat was then placed on its stomach on a slotted aluminum sheet attached to an open plexiglass container. Anesthesia was maintained using a nose cone. A foam pad was placed at the bottom of the plexiglass container. The rat’s head was positioned under a clear plexiglass guide tube where an electromagnet released a 450g steel weight from a 1-meter height. The impact caused the rat to break through the aluminum sheet, undergo a rotational turn of 180 degrees, and land on its back on the foam pad. The weight was attached to a fishing line to prevent a second impact. The nose cone was removed just prior to the weight drop. The rat was removed from the foam pad and returned to its home cage. This produced an acceleration/deceleration injury typical of a mild head injury (Masse et al., 2019). Sham induction followed the same procedure, excluding the impact of weight drop. Rats were randomly assigned to either the weight drop (WtDp) or sham groups (7 sham, 9 WtDp). Rats rested for 24 hours following sham/WtDp induction before returning to daily sessions of the study-test behavioral procedure for a total of 14 sessions post-manipulation. After the sham/WtDp induction, all rats were closely monitored for 24hrs. Rats had no visible injuries (i.e., scarring, skull fractures, bleeding) after the weight drop.

**Figure 3.**
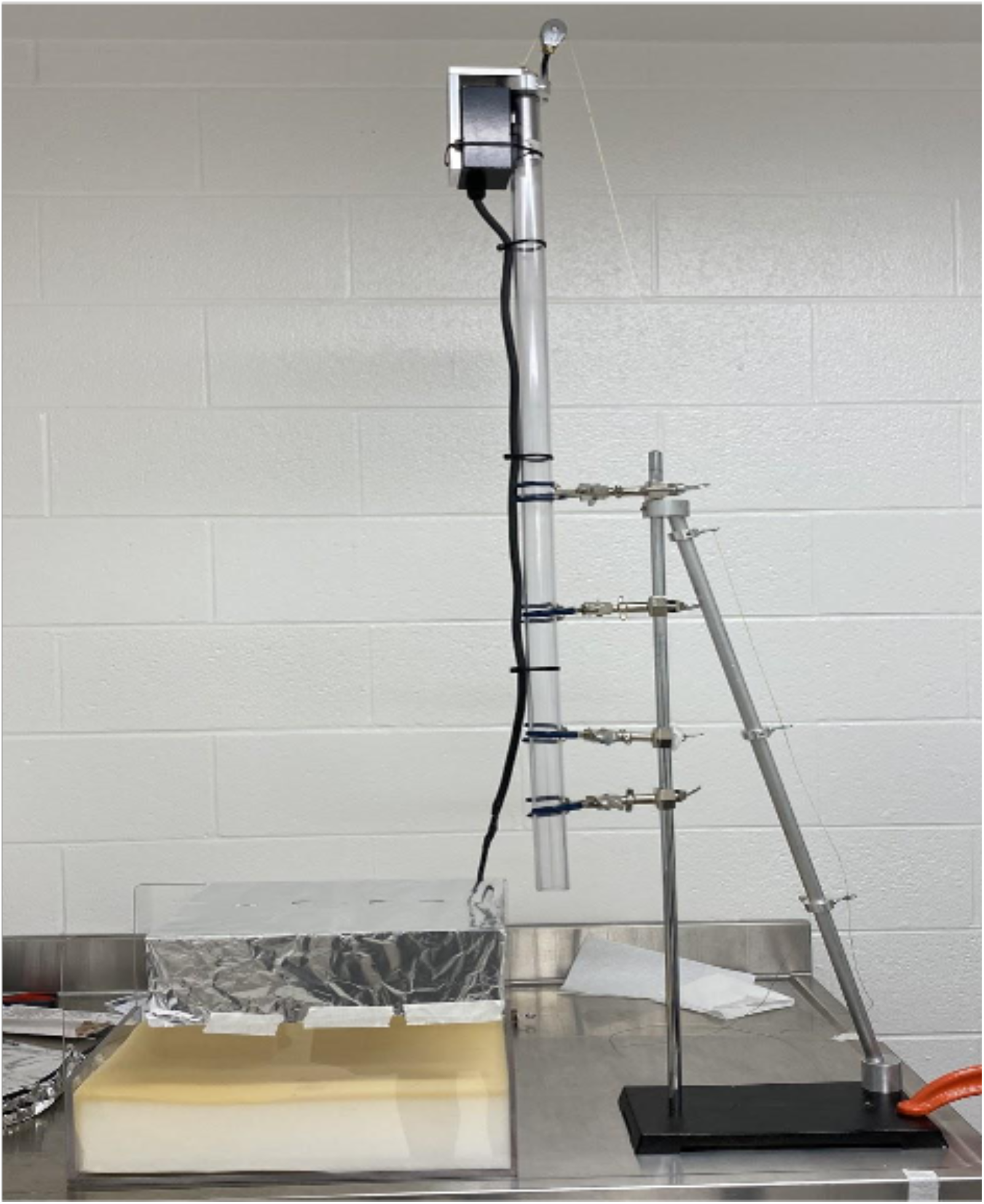
Image of Wayne State University Closed Head Weight Drop Model

## RESULTS

### Behavior

Figure 4 displays, for each group separately, the performance of the animals before and after experimental manipulation. The mean performance is indicated by the red symbol and line. In the sham group (Figure 4A), performance increased from pre- to post-manipulation (mean ± SEM: 7.57 ±3.29, d = 1.2). Five animals showed improved performance from pre- to post-manipulation, whereas two animals showed a decrease. The increase suggests that sham animals continued to learn the task over the post-manipulation sessions. In the WtDp group (Figure 4B), performance declined from pre- to post-manipulation (mean ± SEM: -5.11 ±1.58, d = 0.5). Seven animals show decreased performance from pre- to post-manipulation, whereas two animals show a slight increase. Overall, the weight drop produced a large decline in accuracy relative to the expected increase observed in the sham condition (mean ± SEM: -12.68 ±1.68, d = 1.88). These findings also document medium to large effect sizes in the analysis. There is no qualitative difference between the two sexes for either group. Although the animals were randomly assigned to the two groups, they did not initially perform equally.

**Figure 4.**
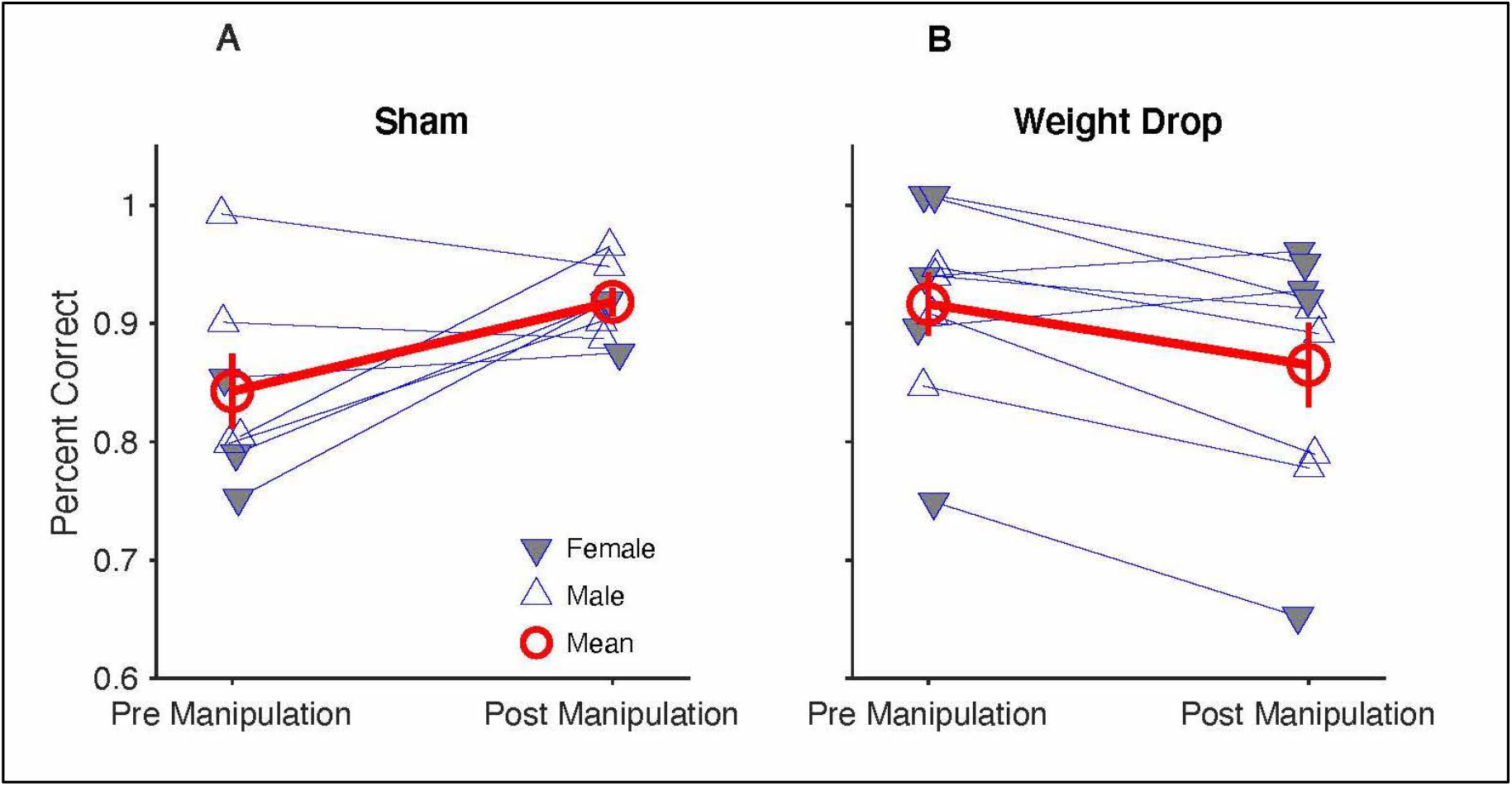
Pre- and post-manipulation performance for the Sham (A) and Weight Drop (B) groups. The mean for each group is shown in red. The Sham group (A) increased in performance from pre- to post-manipulation, while the Weight Drop group (B) declined in performance from pre- to post-manipulation.

Because there was a qualitative difference in performance between groups before manipulation (Sham = 84.3% ±8.3 SEM, Weight Drop = 91.7% ±7.9 SEM), we performed an analysis of covariance (ANCOVA). The model implemented is post-manipulation performance as a function of pre-manipulation performance, sex, and group (sham vs. WtDp). As shown in Table 1, pre-manipulation performance and group performance have a significant effect on post-manipulation performance. There is also a significant interaction between group and pre-manipulation performance, indicating the slopes of post-manipulation performance for the two groups are not parallel. There is no significant effect of sex.

**TABLE 1.**
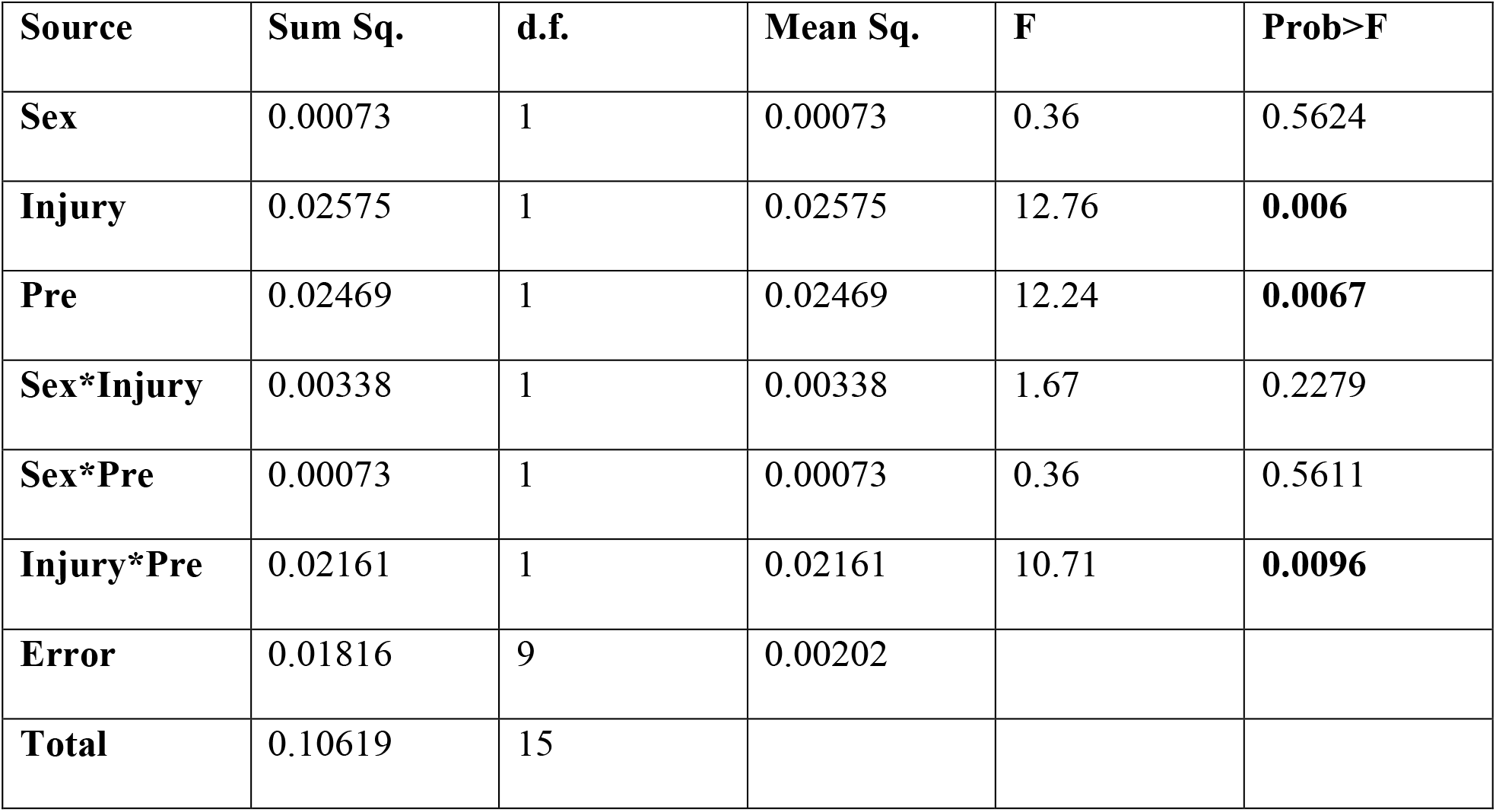
Analysis of Covariance (ANCOVA)

| Source | Sum Sq. | d.f. | Mean Sq. | F | Prob>F |
| --- | --- | --- | --- | --- | --- |
| Sex | 0.00073 | 1 | 0.00073 | 0.36 | 0.5624 |
| Injury | 0.02575 | 1 | 0.02575 | 12.76 | **0.006** |
| Pre | 0.02469 | 1 | 0.02469 | 12.24 | **0.0067** |
| Sex*Injury | 0.00338 | 1 | 0.00338 | 1.67 | 0.2279 |
| Sex*Pre | 0.00073 | 1 | 0.00073 | 0.36 | 0.5611 |
| Injury*Pre | 0.02161 | 1 | 0.02161 | 10.71 | **0.0096** |
| Error | 0.01816 | 9 | 0.00202 |  |  |
| Total | 0.10619 | 15 |  |  |  |

## DISCUSSION

We used a weight drop model to produce an mTBI in rats and examined its effect on spatial working memory. The Wayne State weight drop model produced 12.68% deficit in spatial working memory in the injury group relative to the sham group (d = 1.88).

We made some adjustments to the Wayne State University WDM. The previous model used a hex key to hold the weight in place. Using a hex key could cause variability in how the weight falls because pulling the hex key would shake the tube aimed at the rat’s head. To avoid that, an electromagnet was used to hold the steel weight in place, controlled by an on/off switch, and the tube was clamped to the table to reduce movement.

The results showed rats with a single impact from the Wayne State WDM significantly declined in spatial working memory. In some circumstances, such as sports, people may experience multiple mTBIs over their lifetime. It’s unclear how a repeat injury such as this could affect their memory in the long term. It would be beneficial to determine if this model can be used to examine spatial working memory in a repetitive mTBI paradigm.

Previous studies have shown a sex-related difference in memory impairment (Preiss-Farzanegan, Chapman, Wong, Wu, & Bazarian, 2009). Thus, this study used male and female rats. Researchers are currently using animal models of mTBI to understand better the neurological source that causes those impairments (Kilbourne et al., 2009; Simon-O’Brien, Gauthier, Riban, & Verleye, 2016; Wirth et al., 2017). In this study, male rats were about 150g heavier than the female rats. This difference in weight might influence how each sex is affected by the WDM injury. However, this study did not show any effect of sex on post-manipulation performance, which is similar to what is shown in human mTBI literature (Master et al., 2021).

Memory problems are one of the most common symptoms of mTBI. Previous research has been done using various injury models to assess memory impairment in rats. Rats have been shown to have excellent spatial working memory (e.g. Crystal and Babb 2008). Although the types of injury devices vary, the Wayne State WDM has the potential to closely model human injury. Therefore, our primary goal of this study was to determine if the Wayne State WDM could produce a deficit in spatial working memory, which was achieved. Future studies could also separate the post-manipulation data into acute and chronic timepoints. This would tell us if the impairment of spatial working memory is an acute or a chronic condition.

This study did not evaluate neuroinflammation in the brain. Clinical neuroimaging techniques such as computed tomography (CT) scans and magnetic resonance imaging (MRI) do not detect the subtle changes in the pathology in the brain caused by the damage of mTBI (Ojo, Mouzon, & Crawford, 2016). This makes neuro-pathological studies in humans difficult. For example, the impact of multiple concussions in the development of neuropathology and behavioral impairment is not well characterized.

This knowledge gap also slows down the identification of biomarkers that can confirm brain injury and assist in diagnosing and predicting any lasting effects. This restricted understanding may benefit from the development of animal model research, which may also provide a platform to test pharmaceutical interventions and treatments that may have the ability to reduce or remedy neurological damage. An immunohistochemical analysis should be an additional component of future spatial working memory deficit studies to provide insights into mTBI.

## CONCLUSIONS

Mild traumatic brain injury (concussion) affects millions of people every year. The Wayne State University WDM is a more human-like injury model than the other common injury models. Using this injury model, our study was the first to show a behavioral deficit in spatial working memory in rats. This injury model produced a statistically significant decline in spatial working memory in injured rats relative to sham rats. These research findings are the start of a long journey to understand mTBI and how it affects the brain. Future research should use the Wayne State WDM to determine if it can produce impairment in rats using more complex behavioral tasks.

## ACKNOWLEDGMENTS

This work was supported by supported by NIA AG053524 to JDC

